# Combined Inhibition of SHP2 and CXCR1/2 Promotes Anti-Tumor T Cell Response in NSCLC

**DOI:** 10.1101/2021.03.21.436338

**Authors:** Kwan Ho Tang, Shuai Li, Alireza Khodadadi-Jamayran, Jayu Jen, Han Han, Kayla Guidry, Ting Chen, Yuan Hao, Carmine Fedele, John Zebala, Dean Maeda, James Christensen, Peter Olson, Argus Athanas, Kwok-Kin Wong, Benjamin G. Neel

## Abstract

Clinical trials of SHP2 inhibitors (SHP2i) alone and in various combinations are ongoing for multiple tumors with over-activation of the RAS/ERK pathway. SHP2 plays critical roles in normal cell signaling; hence, SHP2is could influence the tumor microenvironment. We found that SHP2i treatment depleted alveolar and M2-like macrophages and promoted B and T lymphocyte infiltration in *Kras*- and *Egfr*-mutant non-small cell lung cancer (NSCLC). However, treatment also increased intratumor gMDSCs via tumor-intrinsic, NF-kB-dependent production of CXCR2 ligands. Other RAS/ERK pathway inhibitors also induced CXCR2 ligands and gMDSC influx in mice, and CXCR2 ligands were induced in tumors from patients on KRAS^G12C^-inhibitor trials. Combined SHP2(SHP099)/CXCR1/2(SX682) inhibition depleted a specific cluster of *S100a8/9*^high^ gMDSCs, generated *Klrg1*+ CD8+ effector T cells with a strong cytotoxic phenotype but expressing the checkpoint receptor NKG2A, and enhanced survival in *Kras-*and *Egfr*-mutant models. Our results argue for testing RAS/ERK pathway/CXCR1/2/NKG2A inhibitor combinations in NSCLC patients.

**Statement of Significance:** Our study shows that inhibiting the SHP2/RAS/ERK pathway triggers NF-kB-dependent up-regulation of CXCR2 ligands and recruitment of S100A8^high^ gMDSCs, which suppress T cells in NSCLC. Combining SHP2 and CXCR2 inhibitors blocks this gMDSC immigration, resulting in enhanced Th1 polarization, induction of CD8+ KLRG1+ effector T cells with high cytotoxic activity and improved survival in multiple NSCLC models.

## Introduction

SHP2, encoded by *PTPN11*, is required for activation of RAS upstream of the RAS guanine nucleotide exchange proteins SOS1/2. Consequently, SHP2 inhibitors (SHP2i) can block downstream signaling by overactive receptor tyrosine kinases (RTKs) and so-called “cycling” RAS mutants (e.g., KRAS^G12C^), which retain significant intrinsic RAS-GTPase activity and therefore rely on SOS1/2 activity (1). In addition to its potential tumor cell-autonomous actions, SHP2 plays critical roles in normal RTK, cytokine, integrin, and immune checkpoint receptor signaling (2). “Driver” mutations (e.g., amplified or mutant RTKs, mutant KRAS) significantly—and differentially—also have tumor cell-intrinsic and -extrinsic effects and evoke distinct cellular and humoral responses in different tissues (3). Consequently, SHP2is have important, potentially driver-specific, effects on the tumor microenvironment (TME), including potentially complex effects on anti-tumor immunity (2,4-6).

Most pre-clinical studies of SHP2is have used cell-derived or patient-derived xenografts (CDXs, PDXs) established in immune-deficient mice or syngeneic tumor models implanted in the sub-cutaneous (SQ) space. The former models lack adaptive immune responses; the latter rarely harbor the mutational spectrum of the cognate human disease and fail to reproduce tissue-specific immunity (e.g., resident macrophages, T cells, etc.). As the response to targeted therapies in patients almost certainly reflects the composite of direct anti-tumor actions and effects on the TME, CDXs, PDXs, and SQ syngeneic models could provide incomplete or even misleading information about SHP2i action. For example, we found that SHP2i, alone or in combination with KRAS-G12C inhibitor (G12Ci), increased intratumor T cells in KRAS^G12C^-driven non-small cell lung cancer (NSCLC) and pancreas ductal adenocarcinoma (PDAC) (4). However, the degree of T cell function is still unknown. Moreover, combination with anti-PD1 treatment only resulted in minimal improvement in efficacy (4), urging for a more efficacious, rational combination strategy that enhance immune-modulatory effects of SHP2is.

Clinical trials of SHP2is alone or in various combinations are ongoing for multiple disease indications. Systematic characterization of the immune-modulatory effects of SHP2is in genetically-defined, orthotopic or autochthonous, immune-competent tumor models that better reflect human cancers might provide important insights into how best to combine these agents. To this end, we characterized the tumor cell-autonomous and non-autonomous effects of SHP2 inhibition in genetically engineered mouse models (GEMMs) of *Kras*-and *Egfr*-mutant NSCLC and used this information to identify and evaluate a novel, rational combination of SHP2 and CXCR1/2 inhibitors.

## Results

We and others previously demonstrated the efficacy of SHP2i in various *KRAS*-mutant malignancies, including *KRAS*-mutant NSCLC (4,7-11). To systematically explore rational new SHP2i/immune-oncology (I/O) combinations, we first performed a detailed analysis of the effects of the tool compound SHP099 on orthotopic *Kras*^*G12D*^*;Tp53*^*-/-*^ (KP) NSCLC allografts. KP cells were injected intravenously, and lung tumor formation was monitored by magnetic resonance imaging (MRI). Once tumors had reached 100mm^3^ (∼4 weeks) SHP099 (75 mg/kg/d) treatment was initiated. As expected from previous results (4,7), SHP099 had significant single agent efficacy (Fig. 1A). Immune profiling and immunohistochemical staining revealed a significant increase in T lymphocytes in tumor nodules from SHP099-treated mice, compared with vehicle controls without any change in CD8/CD4 ratio (Fig. 1B-C). We also observed a marked reduction in alveolar macrophages (CD11b-CD11c+ Siglec-F+) and M2-like (CD206+/PDL1+) bone marrow (BM)-derived macrophages (CD11b+ CD11c-Ly6C-Ly6G-F4/80+), along with an increase in tumor-infiltrating B lymphocytes (CD19+ cells) (Fig. 1D). Alveolar and M2-like BM-derived macrophages suppress T cell function (12,13), so the observed decrease in these cells could help explain the increased T cell infiltration and anti-tumor effects of SHP099. The effects of B lymphocytes in various tumor models are complex (14-17), although recently, they were been found to exert important anti-tumor effects, including in NSCLC (18,19). Unfortunately, while monocytic myeloid-derived suppressor cells (mMDSC) (CD11b+ Ly6C+ Ly6G-) were unaffected, SHP099 treatment led to a significant increase in infiltrating granulocytic myeloid-derived suppressor cells (gMDSCs) (CD11b+ Ly6G+) (Fig. 1D), consistent with our earlier observation (4). These cells can potently inhibit T cell function (20), and thus could limit the anti-tumor effects of SHP099.

**Figure 1:**
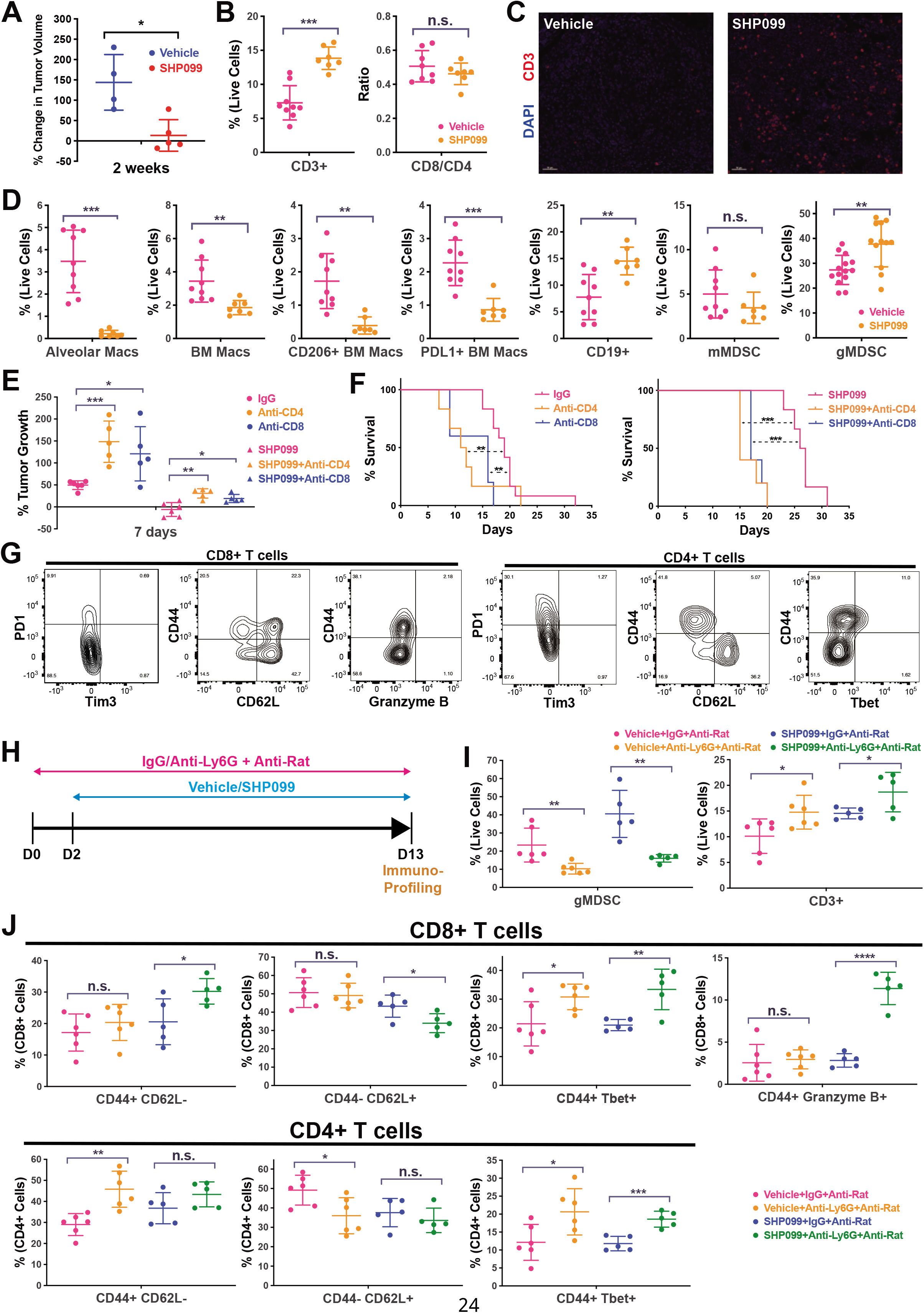
SHP099 treatment promotes T lymphocyte infiltration but not activation. (A) KP NSCLC cells were injected intravenously into C57BL/6 mice. When tumors reached 100mm^3^ (assessed by MRI), treatment was initiated with SHP099 (75mg/kg) or Vehicle daily by gavage. (B) KP allografts were treated with Vehicle or SHP099 for 11 days, and immune profiling was performed on tumor nodules. (C) FFPE sections of representative tumor nodules from mice treated as indicated were stained with an anti-CD3 antibody and visualized with fluorophore-conjugated secondary antibody. (D) Immune profiling of tumor nodules from Vehicle-or SHP099-treated mice after 11 days of treatment. (E) CD4+ or CD8+ T cells were depleted using rat-anti-mouse antibodies (Clones GK1.5 and 2.43 respectively), and tumor growth with/without SHP099 treatment for 7 days was assessed by MRI. (F) CD4+ or CD8+ T cell depletion shortens overall survival of tumor-bearing mice, with/without SHP099 treatment. (G) Phenotypic characterization of infiltrating CD8+ and CD4+ T cell subsets from Vehicle-or SHP099-treated mice (H) Scheme showing gMDSC depletion experiments (I) Depletion of gMDSC promotes greater T cell infiltration, when combined with SHP099 treatment. (J) Effect of gMDSC depletion on phenotypes of infiltrating T lymphocytes. *p< 0.05, **p<0.01, ***p<0.001

We next explored the functional effects of these immune cell populations on KP allograft growth in the absence or presence of SHP099. Depletion experiments indicated significant anti-tumor actions of B cells on KP allografts, as well as secondary effects on specific T cell subsets (Fig. S1A-B and unpublished observations). Mice depleted for CD4 or CD8 T cells also had significantly larger tumors compared with control IgG-injected mice. SHP099 treatment still suppressed KP tumor size, but its effects were reduced in mice lacking either T cell population (Fig. 1E-F). Although SHP099 treatment significantly decreased KP allograft growth, it prolonged median overall survival by only 1 week. Thus, while the infiltrating CD4 and CD8 T cells have demonstrable anti-tumor effects, they clearly cannot orchestrate a sustained anti-tumor response. More detailed characterization of these cells revealed that they did not show a phenotype consistent with “exhaustion” (PD1+ TIM3+), but a significant proportion were naïve (CD44-CD62L+) and only a minority exhibited an effector phenotype (CD44+ CD62L-) (Fig. 1G). Most importantly, only rare infiltrating CD8+ T cells expressed granzyme B (*Gzmb*), a functional marker of cytotoxic T cells (CTL), and only a small fraction of CD4+ T cells expressed Tbet (*Tbx21*), the defining marker for T_H_1 cells (Fig. 1G). These results suggest that although SHP099 treatment promotes T cell immigration into KP tumors, these cells are largely non-functional and have only minimal anti-tumor activity.

We hypothesized that gMDSCs, the largest immune cell population in KP tumors before treatment, whose abundance increases even further following SHP099 administration (Fig. 1D), were responsible for the observed impairment in T cell function. As an initial test of this hypothesis, we depleted gMDSCs by injecting rat anti-Ly6G along with anti-rat antibody (21) and monitored the effects of SHP099 (Fig. 1H). This approach resulted in the expected depletion of gMDSCs along with an increase in intratumor T cells, which increased further in SHP099-treated mice (Fig. 1I). Strikingly, CD8 T cell activation (as marked by increased CD44+/CD62L-and concomitantly decreased CD44-CD62L+ cells) also increased, and there was a marked increase in granzyme B-expressing CD8 cells (Fig. 1J). Depletion of gMDSC also resulted in a basal (without SHP099) increase in activated CD4 T cells and enhanced T_H_1 differentiation in vehicle- and SHP099-treated mice. These results confirm that intratumor gMDSCs exert immune-suppressive effects on tumor-associated CD8+ and CD4+ T cells.

Such antibody-based gMDSC depletion is not accomplished easily in patients. To search for more clinically applicable strategies for preventing SHP2i-induced gMDSC infiltration, we examined immune modulators produced by SHP099-treated KP tumor cells. Transcriptomic analysis identified *Cxcl1* and *Cxcl5*, whose protein products (CXCL1, CXCL5) signal via CXCR2, a major chemotaxis receptor for gMDSC (22), as the most up-regulated chemokines following SHP099 treatment (Fig. 2A, Table S1). *Cxcl1/5* was not induced upon SHP099 treatment of KP cells expressing *PTPN11*^*TM/QL*^, which encodes an SHP099-resistant mutant, as confirmed by pERK immunoblotting (Fig. 2B, C). These results show that *Cxcl1/5* induction is a consequence of SHP2 inhibition, rather than an off-target effect of SHP099. Allografts established with SHP099-resistant KP cells still exhibited significant increases in T and B cell infiltration, along with depletion of alveolar and BM-derived macrophages, following SHP099 treatment. By contrast, they failed to show increased numbers of gMDSCs (Fig. 2D). This result indicates that KP-produced CXCL1/5 is essential for SHP099-evoked gMDSC immigration, emphasizes how SHP2i action reflects complex mix of tumor cell-autonomous and -non-autonomous effects, and comport with a previous report that CXCR2 ligands play important roles in gMDSC recruitment in *KRAS*-mutant colorectal cancer (23).

**Figure 2:**
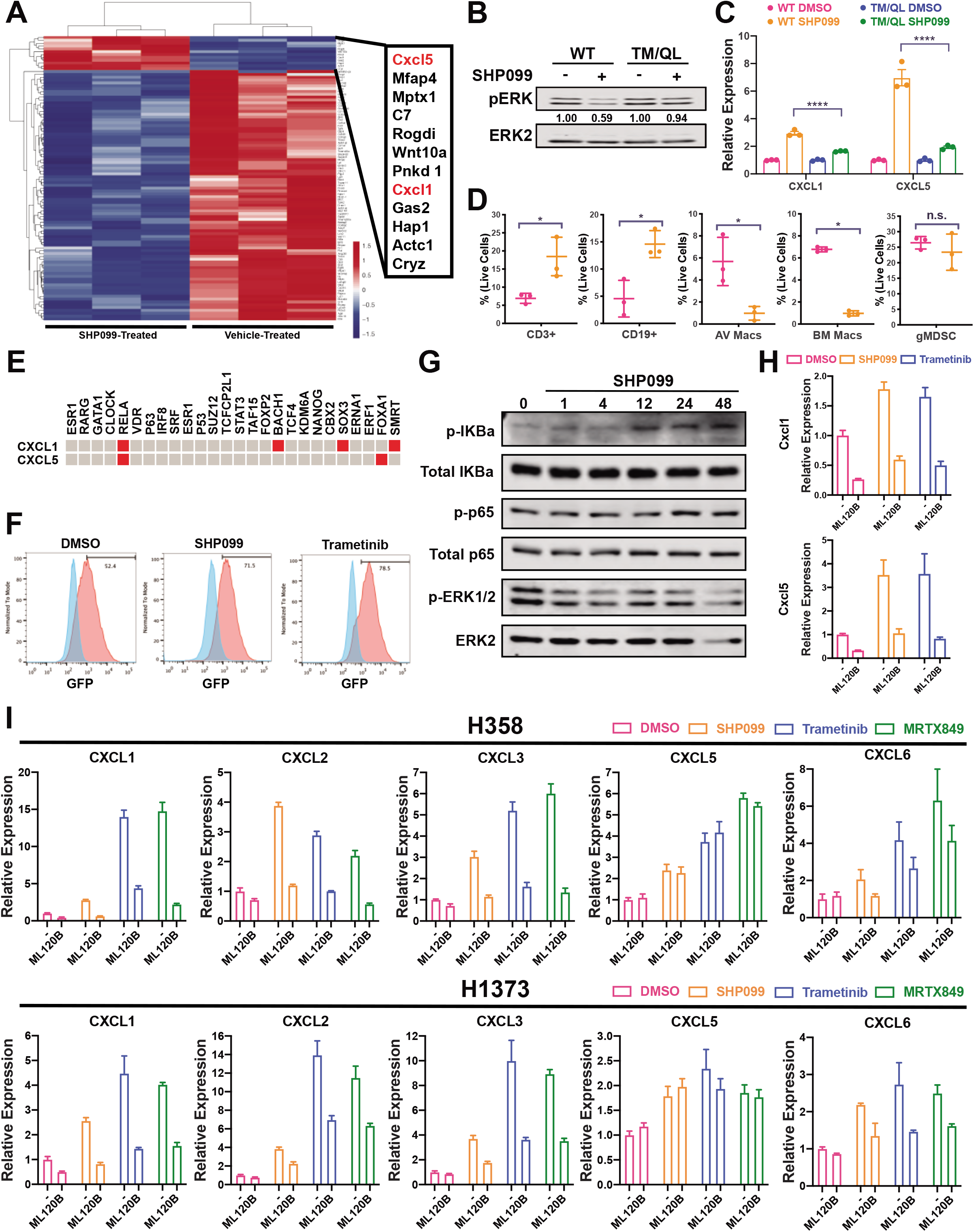
SHP2 inhibition results in NF-kB-evoked *Cxcl1/Cxcl5* up-regulation. (A) RNA-Seq of KP tumors identifies *Cxcl1* and *Cxcl5* as the most up-regulated chemokines after SHP099 treatment. (B) Expressing SHP099-resistant mutant (*PTPN11*^*TM/QL*^) resistant in KP cells prevents SHP099-mediated ERK inhibition. (C) SHP099 treatment significantly induces Cxcl1/5 expression in WT but not SHP099-resistant mutant KP cells. (D) Tumor cell-dependent and -independent effects of SHP099 on immune cells of the tumor microenvironment (E) Promoter-enrichment analysis by Enrichr (GhEA 2016) for *Cxcl1* and *Cxcl5* suggests RELA is active and binds to *Cxcl1* and *Cxcl5* promoters. (F) SHP2 or MEK inhibition increases NF-kB transcriptional activity. An NF-kB-GFP reporter was transduced into KP tumor cells and GFP fluorescence was assessed after 72 hr treatment with SHP099 (10uM), Trametinib (25nM), or DMSO control. (G) Immunoblots showing activation of NF-kB signaling on KP cells treated with 10uM of SHP099. (H) The IKK inhibitor ML120B (10uM) prevents *Cxcl1* and *Cxcl5* up-regulation induced by SHP099 or Trametinib in KP cells (72 hours). (I) Effects of SHP2, KRAS, MEK, and or NF-kB inhibition on GRO family cytokine expression in select human *KRAS*-mutant NSCLC cell lines. *p< 0.05, **p < 0.01, ***p < 0.001

Promoter-enrichment analysis of the KP cell RNA-Seq data suggested activation of genes with RELA binding sites (p=0.012), which include *Cxcl1/5* (Fig. 2E) and *CXCL6* (see below). As RELA is an NF-kB family member, we evaluated the effects of SHP2i on the transcriptional activity of an GFP reporter containing 4 NF-kB sites was stably introduced into KP cells. Consistent with a functional role for NF-KB, SHP099 treatment significantly increased reporter expression as shown by flow cytometry for GFP (Fig. 2F). Treatment of KP cells with the MEK inhibitor Trametinib also evoked increased NF-kB reporter activity and induced *Cxcl1/5* expression (Fig. 2F). These results suggested that NF-kB-mediated *Cxcl1/5* induction might be a general response to RAS/ERK pathway inhibition in KP cells. Immunoblotting showed that, after a significant delay, SHP2 inhibition (and most likely, RAS/ERK pathway inhibition) induced NF-kB pathway activation upstream of I-KB (Fig. 2G). Combining SHP099 or Trametinib with the IKK inhibitor ML120B abolished transcriptional up-regulation of *Cxcl1/5*, confirming the NF-kB-dependence of *Cxcl1/5* induction upon SHP2/MEK inhibition (Fig. 2H). We then explored the potential relevance of these observations to human *KRAS*-mutant NSCLC. CXCL1/5 are members of the GRO family of chemokines, whose genes reside in a common chromosomal region (4q13.3 in humans) and are often co-regulated. Treatment of the *KRAS*^*G12C*^-mutant cell lines H358, H1373 or H2122 with SHP2i-, MEKi-, or the KRAS^G12C^ inhibitor (G12Ci) MRTX849 led to upregulation of multiple GRO family genes, including CXCL1 and CXCL6 (note that *CXCL6* is the homolog of mouse *Cxcl5*). Induction of most of these genes was blocked by ML120B treatment (Fig. 2I, S2A). These results predict that increased immigration of gMDSCs (and perhaps other GRO-dependent immune modulatory cells) might limit the efficacy of SHP2is, MEKis, or G12Cis in human NSCLC as well. Consistent with these observations, we observed increased gMDSC infiltration in tumors from mice treated with Trametinib or MRTX849 (Fig. S2B).

SX682 is a potent inhibitor of CXCR2 (23,24) currently in multiple clinical trials (NCT03161431, NCT04477343, NCT04574583, NCT04245397). Given the results above, we tested the effects of combined SHP2 and CXCR1/2 inhibition. When combined with SHP099, SX682 significantly reduced (although did not eliminate) gMDSC infiltration compared with SHP099 alone (Fig. 3A). Combination therapy also led to a further increase (over single agent SHP099) in intratumor CD4 and CD8 T cells, which exhibited a stronger effector phenotype and enhanced proliferation. The tumor-associated CD8+ T cells in combination-treated groups expressed higher levels of Tbet and granzyme B, consistent with a stronger cytolytic phenotype, while T_H_1 polarization (increased percentage of Tbet+ and decreased percentage of GATA3+ cells) was enhanced in the CD4+ population. Injection of anti-CXCL1 and –CXCL5 neutralizing antibodies had similar effects on gMDSCs and T cells (Fig. S3A-B), providing further evidence that the adverse effects of SHP2i on promoting gMDSC infiltration are driven mainly by CXCL1/5 activation of CXCR2.

**Figure 3:**
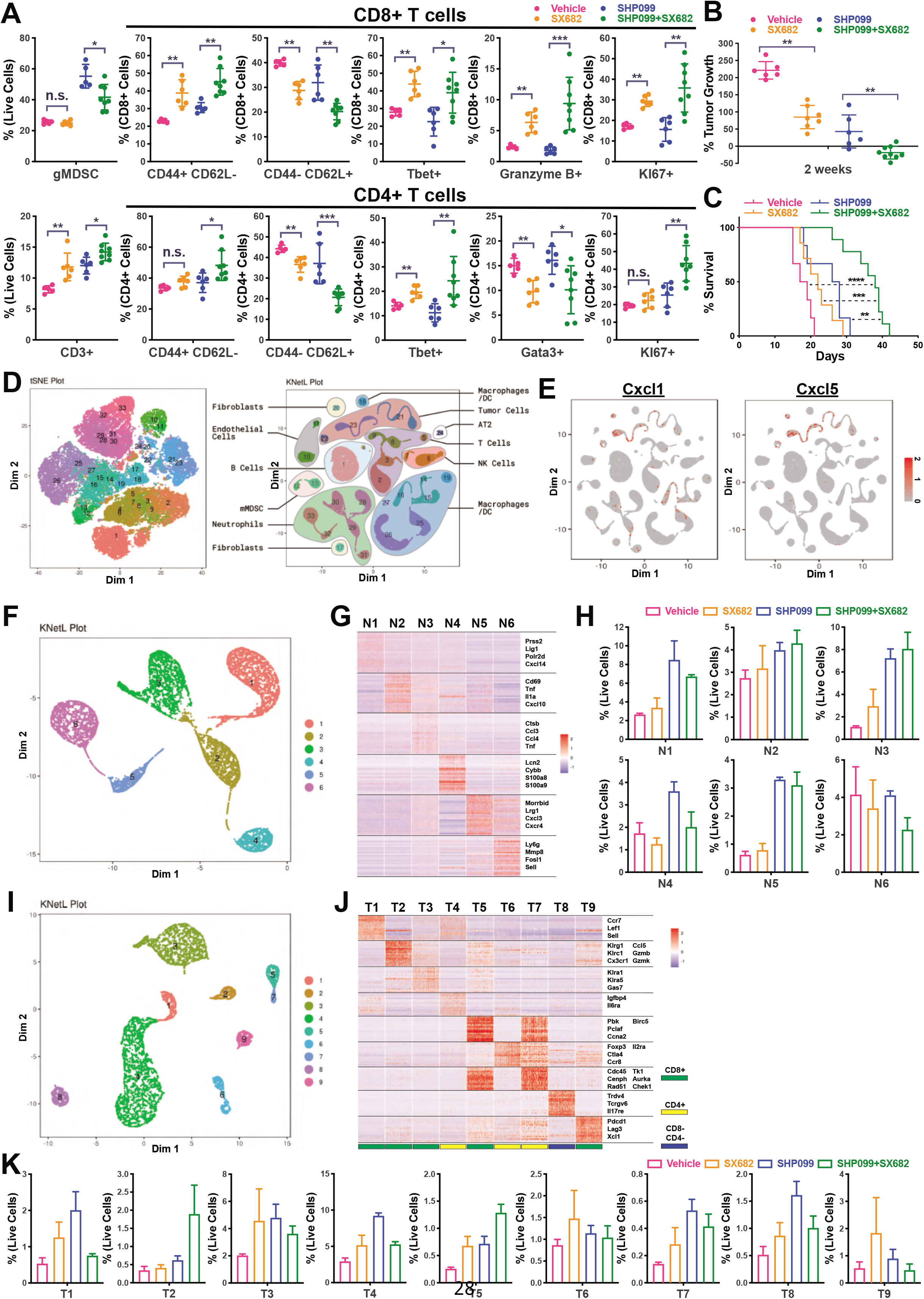
Combined inhibition of SHP2 and CXCR2 promotes anti-tumor T cell response. (A) SX682 significantly suppresses gMDSC infiltration induced by SHP099 and further increases T cell infiltration and proliferation, whereas SX682/SHP099 combination enhances CD8+ and CD4+ T cell activation and induces CD8 CTLs and T_H_1 polarization of CD4 T cells. (B) SX682/SHP099 effectively blocks KP tumor growth at 2 weeks of treatment (as measured by MRI). (C) SX682/SHP099 prolongs overall survival compared with Vehicle-or single agent-treated mice. (D) scRNA-Seq was performed on cells isolated from treated tumors (n = 3 per treatment group) and analyzed by tSNE, or KNetL map. Note the additional clusters revealed by the latter. (E) *Cxcl1* and *Cxcl5* expression are largely restricted to tumor cells and CAFs. (F-G) Identification of six clusters of *S100a8/9*+ gMDSC with distinct transcriptional profiles. (H) SHP099/SX682 specifically depletes cells in cluster N4, which also is induced by SHP099 treatment. (I-J) Identification of 9 clusters of *Cd3e*+ T cells with distinct transcriptional profiles. (K) SHP099/SX682 specifically induces cells in Cluster T2 and T5. *p<0.05, **p< 0.01, ***p < 0.001

Treatment with single agent SHP099 or SX682 for two weeks, at which time vehicle-treated mice start to die, significantly reduced, but did not eliminate, KP allograft growth (Fig. 3B). By contrast, the SHP099/SX682 combination completely suppressed tumor growth at this time point. More importantly, combination treatment significantly prolonged the survival of KP tumor-bearing mice (median: 38 days) as compared to single agent SHP099 (median: 27 days) or SX682 (median: 21.5 days), and more than doubled overall survival, compared with vehicle-treated mice (median: 18 days) (Fig. 3C). There was no evidence of toxicity following long-term (over 5 weeks) SHP099/SX682 combo treatment (Fig. S3C-D and data not shown).

Previous work indicated that CD11b and Ly6G expression (e.g., CD11b+ Ly6G+ cells) alone do not reliably distinguish gMDSCs from normal neutrophils (25,26). To systematically explore the potential heterogeneity in the gMDSC populations following SHP099/SX682 treatment, we performed scRNA-Seq. To analyze the data, we used K-nearest-neighbor-based Network graph drawing Layout (KNetL) map analysis, a graph drawing-based dimensionality reduction algorithm that shows better distinctions in the complex cell communities as compared with tSNE or UMAP analysis (Fig. 3D, S4A). As expected, *Cxcl1* and *Cxcl5* were largely restricted to tumor cells, although there was some expression in cancer-associated fibroblasts (CAFs) (Fig. 3E). Consistent with a previous survey of human NSCLC and KP allograft model (27), we identified six (N1-N6) *S100a8*+ granulocytic cell clusters in KP tumors (Fig. 3F-G, Table S2). Most of these (N1-5) increased after SHP099 treatment, comporting with our flow cytometry results. Interestingly, however (and consistent with the incomplete reduction in the gMDSC population by flow cytometric analysis, Fig. 3A), co-administration of SX682 only blocked infiltration of cells in cluster N4 (Fig. 3H). Notably, N4 cells express significantly higher levels of *S100a8/9*, compared with the other granulocytic cell populations (Fig. 3G), a phenotype that distinguishes gMDSC from normal neutrophils (25,26,28,29). N4 cells also preferentially express *Cybb* (which encodes NOX2) and *Lcn2* (Fig. 3G), raising the possibility that they might suppress T cells by generating superoxide and/or inducing apoptosis in T cells (20,30). N2 and N3 cells are also induced upon SHP099 treatment but were unaffected by co-administration of SX682 (Fig. 3H). N2 and N3 cells preferentially express genes encoding cytokines and chemokines including TNFα, IL1α, CXCL10, CCL3, and CCL4, which promote T cell recruitment and/or T cell activation/differentiation (31,32). These findings suggest a pro-inflammatory role for these granulocytic cells.

Although flow cytometric analysis revealed induction of cytotoxic markers in the CD8+ T cells and more T_H_1 differentiation in combination-treated mice (Fig. 3A), scRNA-Seq also provided insight into the phenotypic heterogeneity of the T cell population. We identified nine clusters (T1-T9) of *Cd3e*+ T cells in KP tumors (Fig. 3I-J, Table S3). SHP099/SX682 treatment led to a specific increase in cells in the *Cd8*+ clusters T2 and T5 (Fig. 3J-K). Cluster T2 comprises *Klrg1*+ *Cx3cr1*+ effector T cells (Teffs) that preferentially express cytotoxic genes including *Gzmb* and *Gzmk* (Fig. 3J). KLRG1 marks highly cytotoxic and proliferative CD8+ Teffs in other settings (33-36). Interestingly, these Teffs also expressed *Klrc1*, which encodes the immune checkpoint receptor NKG2A, but not *Pdcd1, Ctla4* or *Lag3* (Fig. 3J). NKG2A blockade was recently suggested to enhance anti-tumor immunity in subcutaneously syngeneic model (see Discussion) (37). Cells in cluster T5 preferentially express genes associated with proliferation (e.g., *Ccna2, Aurka, Tk1*) but not with effector function (Fig. 3J). In concert, these findings suggest that CXCR2 inhibition specifically blocks infiltration of *S100a8/9*^high^ gMDSCs induced by SHP2i, which in turn lead to the generation of *Klrg1*+ CD8+ Teffs with high cytotoxic and proliferative capability.

Activating mutations in *EGFR* are another major cause of NSCLC, and SHP2 also plays a critical role in EGFR signaling. Given the effects of SHP099 on the KP TME, we asked whether SHP099/SX682 might also have utility in *Egfr*-mutant NSCLC GEMMs. Consistent with a previous report (38), we first confirmed that SHP099 possesses cell-intrinsic ability to suppress MEK-ERK activity and proliferation in *EGFR*-mutant NSCLC cell lines (Fig. 4A-B). SHP2i or MEKi induced NF-kB-dependent expression of several CXCR2 ligands in these lines, Comporting with their effects on *KRAS*-mutant NSCLC cells (Fig. 4C). We then tested SHP099 and SX682 alone or in combination in an osimertinib-resistant *Egfr* mutant GEMM (*Egfr*^*T790M,L858R,C797S*^, hereafter EGFR-TLC). The EGFR driver in this model has the classic L858R mutation, a gatekeeper mutation (T790M) that renders it resistant to first generation EGFR inhibitors, and the C797S mutation that eliminates covalent binding of Osimertinib (Fig. 4D).

**Figure 4:**
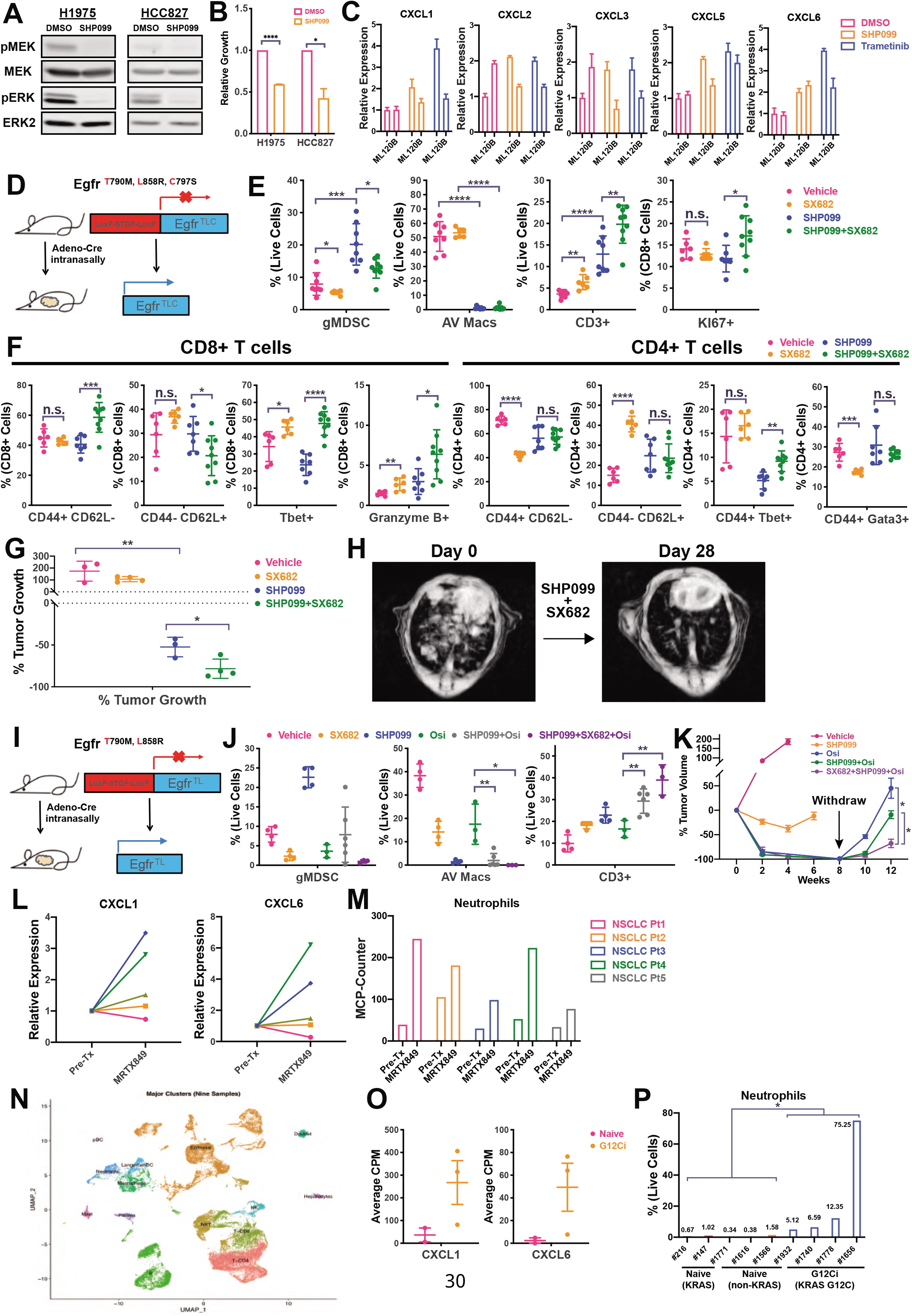
Combined inhibition of SHP2 and CXCR2 also is efficacious in *EGFR-* mutant NSCLC. (A) SHP099 (10uM, 1 hour) inhibits MEK/ERK signaling in human *EGFR*-mutant NSCLC cell lines. (B) SHP099 (10uM, 3 days) suppresses proliferation of human *EGFR*-mutant NSCLC cell lines. (C) SHP099 or Trametinib treatment induces GRO-family chemokine expression in H1975 cells (72 hours treatment), and induction is blocked by IKK inhibitor ML120b. (D) Schematic of Osimertinib-resistant GEMM (EGFR-TLC). (E-F) Effects of SHP099, SX682, and SHP099/SX682 on the indicated immune cell populations in EGFR-TLC tumors. (G-H) SHP099/SX682 significantly enhances anti-tumor efficacy compared with either single agent, quantified after 4 weeks of treatment by MRI (G) and with representative image shown in H. (I) Schematic of Osimertinib-sensitive GEMM (EGFR-TL). (J) Osimertinib/SHP099/SX682 (5mg/kg/, 75mg/kg, 100mg/kg daily) induces significantly stronger T cell response in EGFR-TL GEMM, compared with single agent Osimertinib (5mg/kg) or Osimertinib/SHP099 (5mg/kg, 75mg/kg). (K) Osimertinib/SHP099/SX682 significantly delays disease relapse following drug withdrawal in EGFR-TL mice, compared with mice treated with Osimertinib alone or Osimertinib/SHP099 (doses as above). (L) *CXCL1* and *CXCL6* expression (by RNA-Seq) in paired specimens from NSCLC patients before and after MRTX849 treatment. (L) MCP-Counter analysis of RNA-Seq data reveals increased neutrophil signature in all MRTX849-treated tumors. (M) scRNA-Seq performed on cells isolated from fresh tumor biopsies from NSCLC patients with/without G12Ci treatment. (N) Increased expression of *CXCL1* and *CXCL6* in tumor cells from G12Ci-treated NSCLC patients, compared with those from G12Ci-naïve patients. (O) Significantly higher proportions of neutrophils were observed in tumors from G12Ci-treated patients, compared with those not receiving G12Ci. *p < 0.05, *p<0.01, ***p<0.001

Osimertinib-resistant *EGFR*-mutant NSCLC constitutes a major unmet medical need, as there is currently no effective treatment for these tumors. Compared with KP tumors (Fig. 1B, 1D, EGFR-TLC tumors contained more alveolar macrophages and fewer T cells and gMDSCs (Fig. 4E). Consistent with its effects on KP allografts, SHP099 treatment depleted alveolar macrophages and increased T cell infiltration in tumor-bearing EGFR-TLC mice, but also increased intratumor gMDSCs (Fig. 4E). Single agent SHP099 also failed to evoke T cell activation, CD8+ CTLs or Th1 cells in this *EGFR*-mutant model (Fig. 4F). By contrast, SHP099/SX682 significantly suppressed gMDSC infiltration and promoted more T cell infiltration, greater activation of CD8+ T cells, accompanied by increased proliferation and acquisition of cytolytic phenotype, and enhanced T_H_1 polarization of CD4+ T cells (Fig. 4F). Single agent SHP099 led to a 50% reduction in tumor size by 4 weeks, but SHP099/SX682 significantly enhanced treatment efficacy, leading to >80% tumor shrinkage (Fig. 4G-H).

We also investigated the effects of “up-front” administration of the triple combination of SHP099, SX682, and osimertinib in an osimertinib-sensitive GEMM, *Egfr*^*T790M,L858R*^ (EGFR-TL, Fig. 4I). Again, SHP099 alone or in combination, strongly depleted alveolar macrophages, while increasing intratumor gMDSCs and T cells. The SHP099/SX682/Osimertinib combination resulted in lower levels of gMDSCs and evoked the largest increase in tumor-associated T cells (Fig. 4J). Single agent SHP099 treatment had largely cytostatic effects in this model, while Osimertinib-containing combinations led to complete responses, which were durable for at least 8 weeks (Fig. 4K). Upon drug withdrawal, however, there were marked differences in the rate of tumor recurrence, with single agent Osi-treated mice recurring fastest, followed by the Osi/SHP099 group, and finally mice treated with the three-drug combination (Fig. 4K).

Finally, we asked whether RAS/ERK pathway inhibition results in induction of CXCR2 ligand genes and gMDSC recruitment in NSCLC patients. Remarkably, *CXCL1* and *CXCL6* mRNA levels were increased after MRTX849 treatment in matched biopsy samples from 4/5 *KRAS*^*G12C*^-mutant NSCLC patients (Fig. 4L), while all of the MRTX849-treated patients showed a substantially stronger neutrophil transcriptional signature (Fig. 4M). We also performed scRNA-Seq on several (unmatched) biopsy samples from G12Ci-naïve and G12Ci-treated NSCLC patients (Fig. 4N). As predicted by our pre-clinical studies, tumor cells from G12Ci-treated patients had higher levels of *CXCL1* and *CXCL6* than those from G12Ci-naïve patients (Fig. 4O). We also observed significantly higher proportions of gMDSCs in G12Ci-treated, compared with G12Ci-naïve or non-*KRAS* mutant tumors (Fig. 4P).

## Discussion

SHP2 inhibitors have anti-tumor effects in *KRAS*-mutant and *EGFR*-mutant NSCLC GEMMs, and initial reports demonstrate efficacy of a clinical grade SHP2 inhibitor on patients with cycling KRAS-mutant NSCLC (4-10,38). The extent to which these therapeutic effects reflect direct actions on cancer cells versus cells in the TME has not been studied extensively in orthotopic or autochthonous models. Potentially adverse consequences of SHP2 inhibition, particularly on tumor-associated immune cells, which might limit the efficacy of SHP2is alone or in combination, have not been defined. We find that in addition to direct anti-tumor effects on the highly aggressive KP allograft model, the tool SHP2i SHP099 has several beneficial effects on the immune TME, lowering the level of tumor-promoting alveolar macrophages and M2 BM-derived macrophages, while increasing tumor-associated B and T lymphocytes. The SHP2i-evoked T cells have significant anti-tumor effects as revealed by depletion experiments. However, these effects are limited because SHP099 also induces immigration of gMDSCs, which suppress T cell activation, proliferation, and cytolytic differentiation. Moreover, analysis of pre-and post-treatment biopsies that similar events occur in NSCLC patients on RAS^G12C^ inhibitor trials.

Others have reported that SHP2is evoke meaningful anti-tumor T cell responses (5,6). These studies analyzed subcutaneous syngeneic tumors, which lack tissue-specific immunity and also display mutational spectrums that do not reflect the cognate human malignancies. We previously showed that the same syngeneic cancer cells evoked different TMEs and had quantitatively different drug responses when established in subcutaneous vs. orthotopic sites (4), in accord with other reports (39). Our group previously demonstrated that SHP2i, alone or in combination with G12Ci increased intratumor T cells in in KRAS^G12C^-driven NSCLC and PDAC. Consistent with the effects observed here, most of those T cells failed to express granzyme B (4). Our depletion studies, flow cytometry, and scRNA-Seq analysis reveal that SHP2 inhibition alone, despite evoking significant T cell infiltration in multiple genetically-defined, orthotopic and autochthonous NSCLC models, is unable to generate CD8+ Teff nor enable a strong, highly effective anti-tumor T cell response.

Transcriptional profiling, reporter assays, inhibitor treatment, and neutralizing antibody studies indicate that gMDSC immigration is a consequence of NF-kB-dependent CXCL1/5 production by KP tumor cells. Increased NF-kB-driven transcription appears to result from decreased ERK activation downstream of SHP099, as similar effects are observed upon MEKi and G12Ci treatment. Furthermore, analogous induction of CXCR2 ligands occurs upon SHP2i, G12Ci, or MEKi treatment of human *KRAS*-mutant NSCLC lines. Most importantly, similar induction of CXCR2 ligands and evidence of increased gMDSC infiltration is seen in biopsy samples from patients treated with two different G12Cis. We found that SHP2i or MEKi treatment of *EGFR-*mutant NSCLC lines also induced such chemokines. Taken together, these data suggest that CXCR2 ligand induction (and possibly gMDSC immigration) might be a general response to SHP2i/MEKi treatment.

Earlier studies assigned tumor-infiltrating granulocytes to “anti-tumor” N1 or “pro-tumor” N2 subsets (31). N1 granulocytes secrete pro-inflammatory cytokines (e.g., TNFα, CXCL10, IL12) that facilitate anti-tumor T cell responses; N2 subsets suppress or kill T cells via production of reactive oxygen species (ROS), arginase, and/or nitric oxide (20,31). A previous survey of a large number of human and mouse lung tumors by scRNA-Seq revealed more extensive phenotypic heterogeneity (27), but the functional importance of these subpopulations remains largely unknown. Recent reports suggest that gMDSCs can be distinguished from normal neutrophils by virtue of high S100a8/9 expression (25,26,28,29). Consistent with these findings, we identified 6 unique subsets of tumor-infiltrating granulocytes, with transcriptomic profiles suggesting pro-inflammatory (N2: *Tnf, Il1a, Cxcl10*; N3: *Ccl3, Ccl4, Tnf*) and immuno-suppressive (N4: *S100a8/9, Cybb, Lcn2*) roles, respectively. SHP099 treatment evokes increased infiltration of nearly all granulocytic sub-populations into KP tumors, including those with pro-inflammatory and immuno-suppressive function. Intriguingly, co-administration of SX682 specifically blocks infiltration of *S100a8/9*^high^ immuno-suppressive (N4) granulocytes. NOX2 is an NADPH oxidase, which catalyzes superoxide generation, whereas LCN2 has iron chelating function and reportedly induces T cell apoptosis (30). S100A8/9 secreted by gMDSC also can re-activate dormant KP tumor cells (28). Dormancy is increasingly recognized as a key mechanism by which tumor cells, including *KRAS*-mutant NSCLC (40), escape the effects of targeted therapy and even conventional chemotherapy (41-43). Hence, blocking N4/S100A8/9^high^ gMDSC infiltration by CXCR2 inhibition could have multiple beneficial effects in therapeutic combinations. Because CXCR1/2 inhibition specifically blocks immuno-suppressive gMDSCs without impairing pro-inflammatory granulocyte infiltration, and SX682 and other CXCR1/2 inhibitors already are in clinical trials (NCT03177187, NCT03473925), our findings could have immediate implications for the clinic.

Consistent with the above analysis, gMDSC-mediated suppression by SX682 treatment resulted in increased numbers of intratumor CD8 T cells. These cells are more activated and exhibit *Klrg1*+ Teff phenotype marked by high expression of cytolytic genes. Intratumor CD4 T cell number, activation, and T_H_1 polarization also increased. Notably, KLRG1 marks a subset of highly cytotoxic, proliferative, and often short-lived CD8+ Teffs induced in the setting of certain infections by the combination of strong TCR and inflammatory signals (33-36). For example, *Klrg*^-/-^ mice have more total and activated CD4+ T cells and survive longer after infection with *Mycobacterium tuberculosis* (44), suggesting that KLRG might function as an immune checkpoint receptor. However, its role in cancer remains largely unknown.

Intriguingly, we found that these CD8+ *Klrg1*+ Teffs preferentially express the checkpoint *Klrc1* (NKG2A) but not conventional checkpoint receptors such as PD1, CTLA4, and LAG3. Such findings may explain why *KRAS*-mutant NSCLC responds incompletely to PD1/PD-L1 blockade (45,46). NKG2A is best known as a Killer Inhibitory Receptor (KIR) on NK cells (47,48). However, more recent work shows that it also is expressed in early CD8+ Teffs, and NKG2A blockade significantly enhances anti-tumor protective immunity induced by cancer vaccines (37). Anti-NKG2A mAb (Monalizumab) treatment also was found to enhance CD8+ T cell function in a phase 2 clinical trial (49), and a phase 3 trial of the effects of NKG2A blockade is undergoing (NCT04590963).

Notably, combined SHP099/SX682 inhibition more than doubled the survival of mice bearing extremely aggressive KP allografts and substantially reduced tumor burden in Osimertinib-sensitive and -resistant NSCLC GEMMs. Although our results support the testing of SHP2i/CXCR1/2 combinations in patients, our analysis of the remaining T cell populations suggests that combining SHP2i (and/or other RAS/ERK pathway inhibitors) with CXCR2 and NKG2A blockade might be even more efficacious. As drugs targeting each are currently be tested in various clinical trials, expeditious testing these concepts should be possible. While our focus here was on *KRAS-*and *EGFR-*mutant NSCLC, our results suggest that induction of CXCR2 ligands and consequently, increased granulocytic infiltration into tumors could be a common, unavoidable “side-effect” of RAS/ERK pathway inhibition in other NSCLC subsets and tumor types. Finally, it will be interesting to see if normal cells (e.g., lung, GI, liver) also produce CXCR2 ligands in response to RAS/ERK pathway inhibition. If so, then granulocytic infiltration as a consequence of these ligands might contribute to the well-known inflammatory side effects of these drugs.

## Acknowledgement

We thank the PCC Experimental Pathology, Precision Immunology, Genome Technology Center, and Applied Bioinformatics Laboratories for technical support. We thank Drs. Toshiyuki Araki, Kiyomi Araki, Abhishek Bhardwaj, Mitchell Geer, Connor Foster, Wei Wei, Jiehui Deng for helpful advice and discussions. Carmine Fedele is currently affiliated with Novartis Institutes for Biomedical Research. This work was supported by National Institutes of Health grants CA49152 (B.G. Neel), CA248896 (B.G. Neel and K.K. Wong) and Cancer Center Core Grant P30 CA016087.

## Methods

### Cell Culture and Reagents

Mouse KP tumor cells, H358, H1373, H2122, H23, and H1975 were from stocks in the Wong laboratory. Cells were cultured in RPMI media supplemented with 10% fetal bovine serum and 1% Penicillin-Streptomycin in a 37°C incubator with 5% CO2. Cells were tested routinely (every 3 months) for mycoplasma contamination by PCR (Young et al., 2010), and genotyped by STR analysis at IDEXX Bioresearch. SHP099 was synthesized by Wuxi AppTec. SX682 was obtained from Syntrix Pharmaceuticals. Trametinib was purchased from Selleckchem. The NF-kB-GFP-Reporter was purchased from System Biosciences. The *PTPN11*^T253M/Q257L^ (TM/QL) expression construct was reported previously (50).

### Lentivirus Production

Viruses were produced by co-transfecting HEK293T cells with lentiviral constructs and packaging vectors (pVSV-G+dR8.91) in DMEM supplemented with 10% fetal bovine serum. Transfection media were replaced by fresh media after 12 hours. Virus-containing supernatants were collected 60 hours later, passed through 0.45um filters, and then used to infect various cultured cells in the presence of 8ug/mL polybrene (Sigma).

### Animal Studies

All animal studies were reviewed and approved by the Institutional Animal Care and Use Committee (IACUC) at New York University Grossman School of Medicine. For the orthotopic allograft lung cancer model, six-week-old male B6 WT mice were purchased from The Jackson Laboratories. KP cells (10^6^) in 200µL PBS were injected into the tail vein of each mouse. MRI was used to monitor tumor formation and progression in orthotopic models. Mice were treated with vehicle or 75mg/kg SHP099 once daily by oral gavage. For CXCR1/2 inhibitor combination studies, mice were dosed with SX-682 (100 mg/kg, daily), either alone or in combination with SHP099 (75mg/kg daily).

The *EGFR*-mutant NSCLC GEMMs harbor a conditional activating mutation of human of EGFR-L858R/T790M with/without C797S (TLC GEMM or TL GEMM, respectively) at the collagen I locus (51). Cre-recombinase was induced through intranasal inhalation of 5×10^7^p.f.u. adeno-Cre (University of Iowa adenoviral core), and tumors (adenocarcinomas) typically appeared 16 weeks after induction. For drug treatment studies, age-matched littermates (6-8-week-old) were induced, and tumor burden was monitored by MRI. Once tumor size reached 300-400 mm^3^ (∼20 weeks after adenoviral inoculation), mice were randomly assigned to experimental groups. No gender differences were observed in tumor growth or drug response. Mice were evaluated by MRI every other week to quantify lung tumor burden before and after drug treatment. TLC tumor-bearing mice were treated with vehicle, SHP099 (75 mg/kg QD), SX-682 (100mg/kg QD) or both drugs. For TL mice, in addition to these 4 arms, Osimertinib (5mg/kg) was introduced in combination with SHP099 or SHP099+SX-682.

To specifically deplete T cells, 400ug of rat IgG2b (Clone LTF-2, Bioxcell), anti-mouse CD4 antibody (Clone GK1.5, Bioxcell) or anti-mouse CD8 antibody (Clone 2.43, Bioxcell) were injected into the mouse peritoneum (IP) one day before commencement of Vehicle/SHP099 treatment. Subsequently, antibodies (200ug) were administered twice a week throughout the experiments. For specific depletion of gMDSCs, 25ug of rat IgG2a (Clone 2A3, Bioxcell) or anti-mouse Ly6G antibody (Clone 1A8, Bioxcell) were administered daily in the first week, 2 days ahead of the Vehicle/SHP099 treatment. The dose was increased to 50ug starting from the second week; in addition, mice received 50ug of anti-rat secondary antibody (Clone MAR18.5, Bioxcell) every other day. For *in vivo* neutralization of CXCL1 and CXCL5, 80ug of anti-CXCL1 (Clone 20326, Leinco) and anti-CXCL5 (Clone 61905, Leinco) antibodies were administered IP every 5 days. For depletion of B cells, 250µg of anti-CD20 antibody (Clone SA271G2, Biolegend) was administered two days before commencement of Vehicle/SHP099 treatment.

### Magnetic Resonance Imaging

Animals were anesthetized with isoflurane, and magnetic resonance imaging (MRI) of the lung field was performed using the BioSpec USR70/30 horizontal bore system (Bruker) to scan 24 consecutive sections. Overall tumor volumes within the whole lung were quantified using 3D slicer software to reconstruct MRI volumetric measurements as described (52). Acquisition of the MRI signal was adapted according to cardiac and respiratory cycles to minimize motion effects during imaging.

### Cell Isolation

Tumor nodules were resected from lungs with visible tumors. Nodules were then cut into small pieces and digested with Collagenase/Hyaluronidase (StemCell Technologies) and DNase I (StemCell Technologies) in Advance DMEM/F12 media (Gibco) for 45 mins in 37°C. Cell suspensions were then filtered through 70um cell strainers (Fisher) and washed with cold FACS buffer (2% FBS in PBS). Red blood cells were lysed by resuspending cell pellets in ACK Lysis Buffer (Gibco) for 2 mins, followed by washing with cold FACS buffer.

### Flow cytometry

Freshly prepared cell suspensions were blocked with 1% mouse serum (Jackson ImmunoResearch), 1% rat serum (Jackson ImmunoResearch), and 2% mouse FcR Blocking Reagent (Miltenyl) for 15 mins at 4°C. Fluorophore-conjugated primary antibodies against cell surface antigens were added to the cell suspensions and incubated for 30 mins at 4°C. Cells were then washed with cold PBS and stained with LIVE/DEAD UV (Invitrogen) according to manufacturer’s instructions. Cells were then washed with cold FACS buffer and fixed with Foxp3/Transcription Factor Staining Buffer Set (eBioscience) according to manufacturer’s protocol. Fixed cells were again blocked with 1% mouse serum, 1% rat serum, and 2% mouse FcR Blocking Reagent for 15 mins at 4°C and then stained with fluorophore-conjugated primary antibodies against intracellular antigens. Cells were then washed, and data were acquired on a LSR II Flow Analyzer (BD). Details on the antibodies used are provided in Supplementary Table S4. Data acquired were analyzed by using FlowJo software (BD).

### Immunofluorescence

Mice with tumor-bearing lungs were perfused with PBS, fixed in 10% formalin for 48 hours, washed in 70% ethanol, embedded, and sectioned (10um) for histological analysis. Immunofluorescence was performed with the OPAL multiplexed immunofluorescence staining platform by the Experimental Pathology Shared Resource at Perlmutter Cancer Center (PCC).

### RNA Extraction, cDNA Synthesis, and qPCR

Isolated tumor cells or trypsinized cancer cell lines were washed with PBS, and total RNA was extracted from cell pellets by using the miRNeasy Mini Kit (Qiagen). cDNAs were generated by using the SuperScript IV First Strand Synthesis System (Invitrogen). qRT-PCR was performed with Fast SYBR Green Master Mix (Applied Biosystems), following the manufacturer’s protocol, in 384-well format in a C1000 Touch Thermal Cycler (Bio-Rad). Differential gene-expression analysis was performed with CFX Manager (Bio-Rad) and normalized to GAPDH expression. Primers used are listed in Supplementary Table S5.

### Bulk RNA-Seq

RNA-Sequencing was performed on total RNA from isolated tumor cells by the PCC Genome Technology Center Shared Resource (GTC). Libraries were prepared using the Illumina TruSeq Stranded Total RNA Sample Preparation Kit and sequenced on an Illumina NovaSeq 6000 machine using 150bp paired-end reads. Sequencing results were de-multiplexed and converted to FASTQ format using Illumina bcl2fastq software. Subsequent data processing and analysis were performed by the PCC Applied Bioinformatics Laboratories (ABL). Promoter-enrichment analysis was performed on bulk RNA-Seq data by using Enrichr (53,54).

### Single Cell RNA-Seq and Data Analysis

Single cell suspensions isolated from treated tumors are individually barcoded with hashtag antibodies (Anti-mouse TotalSeq-C antibodies, Biolegend). Each sample was washed 4 times with PBS + 2% BSA before pooling. Three replicates from same treatment groups were pooled as one sample (52,000 cells each sample). Specimens were then filtered through 70uM strainers (Fisher). Cell concentration, singularity, and viability were confirmed with a hematocytometer before submission for scRNA-Seq (10X Genomics). Experiments were performed with DNA LoBind 1.5 mL tubes (Eppendorf). scRNA-Seq was performed by the GTC, with subsequent data processing and analysis performed by the ABL. Quality controls included calculation of the number of genes, UMIs, and the proportion of mitochondrial genes for each cell. Cells with a low number of covered genes (gene-count < 500) or high mitochondrial counts (mt-genes > 0.2) were filtered out, and the matrix was normalized based on library size. A general statistical test was performed to calculate gene dispersion, base mean and cell coverage. Genes with high coverage (top 500) and high dispersion (dispersion > 1.5) were chosen to construct the gene model (1890 genes). The iCellR R package (v1.5.5) (https://CRAN.R-project.org/package=iCellR) was used to perform Principal Component Analysis (PCA) and batch alignment on this model. T-distributed Stochastic Neighbor Embedding (t-SNE), Uniform Manifold Approximation and Projection (UMAP) and K-nearest-neighbor-based Network graph drawing Layout (KNetL) were then performed. KNetL map has a zoom option which allows users to see more or less detail (more or fewer sub-populations in cell communities); in the studies here, we used a zoom of 650. The network layout used in KNetL map is force-based (55), and the zoom option changes the force in the system. Force-directed graph drawing algorithms assign attractive (analogous to spring force) and repulsive forces (usually described as analogous to the forces in atomic particles) to separate all pairs of nodes. The network analysis used in KNetL map has long been used for single cell analysis and clustering (56); here, the nodes of the network layout are extracted and UMAP is performed to create the final plot (“KNetL map”). PhenoGraph (56) clustering was then performed on the KNetL map results, and marker genes were found for each cluster and visualized on heatmaps, bar plots, and boxplots as indicate. Marker genes were then used to assign cell types. Imputation was used for some data visualizations only and not for the analysis. For imputation we used KNN to average the expression of 10 neighboring cells per cell, using iCellR’s “run.impute” function on KNetL data.

### Statistical Analysis

Data are expressed as mean ± standard error of the mean. Statistical significance was determined using Student t test or Mann–Whitney U test, as indicated/ Statistical analyses were performed in Prism 9 (GraphPad Software). Significance was set at p=0.05.

---

**Supplementary Table 4.** Antibody list:

| **Antigen** | **Clone** |
| --- | --- |
| CD45 (mouse) | 30-F11 (BD) |
| CD3e (mouse) | 145-2C11 (Biolegend) |
| CD3 (mouse) | CD3-12 (Biorad) |
| CD4 (mouse) | RM4-5 (BD) |
| CD8a (mouse) | 53-6.7 (Biolegend) |
| CD44 (mouse) | IM7 (Biolegend) |
| CD62L (mouse) | MEL-14 (Biolegend) |
| PD1 (mouse) | 29F.1A12 (Biolegend) |
| Tim3 (mouse) | RMT3-23 (Biolegend) |
| Foxp3 (mouse) | FJK-16s (Invitrogen) |
| Tbet (mouse) | 4B10 (Biolegend) |
| Gata3 (mouse) | L50-823 (BD) |
| RORgt (mouse) | Q31-378 (BD) |
| Bcl6 (mouse) | K112-91 (BD) |
| Granzyme B (mouse) | GB11 (BD) |
| Kl67 (mouse) | SolA15 (Invitrogen) |
| CD19 (mouse) | 6D5 (Biolegend) |
| CD11b (mouse) | M1/70 (Biolegend) |
| CD11c (mouse) | HL3 (BD) |
| Siglec-F (mouse) | E50-2440 (BD) |
| CD103 (mouse) | 2E7 (Biolegend) |
| Ly6C (mouse) | HK1.4 (Biolegend) |
| Ly6G (mouse) | 1A8 (Biolegend) |
| F4/80 (mouse) | BM8 (Biolegend) |
| MHC II (mouse) | M5/114.15.2 (Biolegend) |
| iNOS (mouse) | CXNFT (Invitrogen) |
| CD206 (mouse) | C068C2 (Biolegend) |
| PDL1 (mouse) | 10F.9G2 (Biolegend) |

**Supplementary Table 5.**
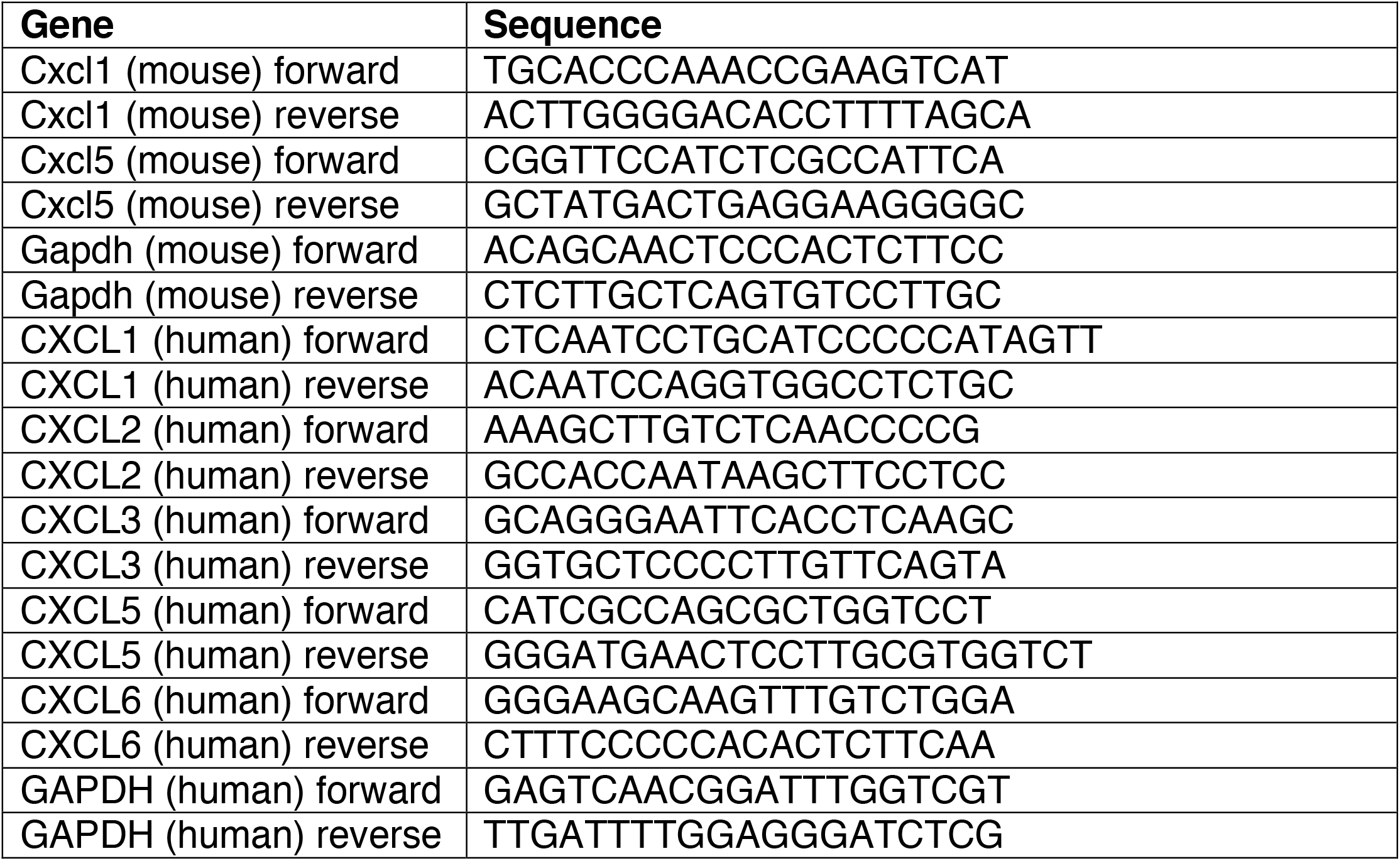
Primer List.

